# Informal genomic surveillance of regional distribution of *Salmonella* Typhi genotypes and antimicrobial resistance via returning travellers

**DOI:** 10.1101/510461

**Authors:** Danielle J. Ingle, Satheesh Nair, Hassan Hartman, Philip M. Ashton, Zoe A. Dyson, Martin Day, Joanne Freedman, Marie A. Chattaway, Kathryn E. Holt, Timothy J. Dallman

## Abstract

*Salmonella enterica* serovar Typhi (*S.* Typhi) is the causative agent of typhoid fever, a systemic human infection with a burden exceeding 20 million cases each year that occur disproportionately among children in low and middle income countries. Antimicrobial therapy is the mainstay for treatment, but resistance to multiple agents is common. Here we report genotypes and antimicrobial resistance (AMR) determinants detected from routine whole-genome sequencing (WGS) of 533 *S*. Typhi isolates referred to Public Health England between April 2014 and April 2017, 488 (92%) of which had accompanying patient travel information obtained via an enhanced surveillance questionnaire. The majority of cases involved *S*. Typhi 4.3.1 (H58) linked with travel to South Asia (58%). Travel to East and West Africa were associated with genotypes 4.3.1 and 3.3.1, respectively. Point mutations in the quinolone resistance determining region (QRDR), associated with reduced susceptibility to fluoroquinolones, were very common (85% of all cases) but the frequency varied significantly by region of travel: 95% in South Asia, 43% in East Africa, 27% in West Africa. QRDR triple mutants, resistant to ciprofloxacin, were restricted to 4.3.1 lineage II and associated with travel to India, accounting for 23% of cases reporting travel to the country. Overall 24% of isolates were MDR, however the frequency varied significantly by region and country of travel: 27% in West Africa, 52% in East Africa, 55% in Pakistan, 24% in Bangladesh, 3% in India. MDR determinants were plasmid-borne (IncHI1 PST2 plasmids) in *S*. Typhi 3.1.1 linked to West Africa, but in all other regions MDR was chromosomally integrated in 4.3.1 lineage I. We propose that routine WGS data from travel-associated cases in industrialised countries could serve as informal sentinel AMR genomic surveillance data for countries where WGS is not available or routinely performed.

**Author Summary:** Our data demonstrate how routine WGS data produced by Public Health England can be further mined for informal passive surveillance of *Salmonella* Typhi circulating in different geographical regions where typhoid is endemic. We have shown the public health utility of a simplified approach to WGS reporting based on the GenoTyphi genotyping framework and nomenclature, which doesn’t require the generation of a phylogenetic tree or other phylogenetic analysis. These approaches yielded results consistent with previously reported antimicrobial resistance (AMR) patterns of *S*. Typhi, including prevalence of multi-drug resistant (MDR) and fluoroquinolone resistance in different regions in association with different pathogen variants. These data provide a rationale and framework for the extraction and reporting of geographically stratified genotype and AMR data from public health labs in non-endemic countries. Prospective analysis and reporting of such data could potentially detect shifts in regional *S*. Typhi populations, such as replacement or spread of different subclades and the emergence and dissemination of MDR, fluoroquinolone resistant and/or extensively drug resistant *S*. Typhi, providing valuable data to inform typhoid control measures in low and middle income countries that are still building their genomics capacity.

## Introduction

*Salmonella enterica* serovar Typhi (*S*. Typhi) is the causative agent of typhoid fever[1], a systemic human infection responsible for an estimated 223,000 deaths each year resulting from more than 20 million infections[2,3]. The majority of the disease burden falls on children in low and middle income countries (LMICs)[2], however vaccination programmes are rare in endemic countries and antimicrobial therapy is considered crucial for the safe clearance of *S*. Typhi infections and the avoidance of clinical complications. Historically typhoid fever could be effectively treated using first-line drugs including chloramphenicol, ampillicin or co-trimoxazole, however the emergence of multi-drug resistant (MDR) *S*. Typhi, defined as displaying resistance to all three drugs, in the late 1980s and early 1990s resulted in changes to treatment guidelines, with fluoroquinolones becoming the recommended therapy[4-7]. Recent years have seen a rise in the proportion of *S*. Typhi disease isolates displaying reduced susceptibility to fluoroquinolones associated with point mutations in quinolone resistance determining region (QRDR) of *gyrA* and parC[2,8,9].

*S*. Typhi is genetically monomorphic, which has historically constrained molecular surveillance of *S*. Typhi before the era of high throughput whole genome sequencing (WGS)[5,10]. A phylogenetically informative genotyping scheme, GenoTyphi, was recently introduced to facilitate the interpretation of *S*. Typhi WGS data[11]. Application of the scheme to a global collection of isolates from >60 countries showed that the *S*. Typhi population is highly structured, comprising dozens of subclades associated with specific geographical regions[11,12]. This global genomic framework for *S*. Typhi revealed the majority of MDR *S*. Typhi infections worldwide are associated with a single phylogenetic lineage, designated as genotype 4.3.1 (H58 under the legacy scheme[5]), which has disseminated from South Asia since the 1990s, including into East Africa [11,12]. Contrastingly, in West Africa MDR *S*. Typhi is associated with a different genotype, 3.1.1[13,14]. The MDR phenotype in both *S*. Typhi 4.3.1 and 3.1.1 is encoded by a composite transposon carrying genes conferring resistance to five drug classes. These include chloramphenicol, penicillins and co-trimoxazole and the determinants are typically located on IncH1 plasmids (plasmid sequence type PST6 in 4.3.1, and PST2 in 3.1.1) [4,12-17]. Recently, migration of the MDR transposon to the *S*. Typhi chromosome via IS*1* transposition has been reported, with four different integration sites identified in the *S*. Typhi 4.3.1 strain background (*fbp, yidA,* STY4438 and the intergenic region between *cyaA* and *cyaY*)[12,18-21].

Resistance to newer agents is also on the rise in *S*. Typhi. Reduced susceptibility to fluoroquinolones via QRDR mutations is commonly observed amongst *S*. Typhi 4.3.1[12], especially lineage II[16,22], but is rare in 3.1.1 and other genotypes[12,13]. *S*. Typhi 4.3.1 QRDR triple mutants (carrying two mutations in *gyrA* and one in *parC)* displaying complete resistance to ciprofloxacin are increasing in prevalence in South Asia[23]. WGS analysis showed that recent treatment failure with gatifloxacin in Nepal resulted from introduction of a 4.3.1 lineage II triple mutant from India, prompting a reconsideration of the reliance on fluoroquinolones in the region[8,22]. Subsequently an outbreak of *S*. Typhi 4.3.1 displaying resistance to ceftriaxone in addition to ciprofloxacin and all three first line drugs was reported in Pakistan[24,25]. The outbreak strain carried both the MDR composite transposon integrated in the chromosome at *yidA,* and an *E. coli* IncY plasmid harbouring the extended spectrum beta-lactamase (ESBL) gene *bla_CTX-M-15_* (conferring resistance to ceftriaxone and other third generation cephalosporins) and the quinolone resistance gene *qnrS* (which combined with a gyrA-S83F mutation in the chromosome conferred resistance to ciprofloxacin)[24]. This strain has been designated XDR (extensively drug resistant, defined as resistance to chloramphenicol, penicillins, co-trimoxazole, ceftriaxone and ciprofloxacin) and severely limits treatment options, with azithromycin being the last remaining oral antibiotic (to which sporadic resistance has already been observed in South Asia)[7].

These dynamic trends highlight a need for prospective AMR surveillance in global *S*. Typhi populations, in order to inform empirical treatment options; genomic surveillance offers the added benefit of revealing resistance mechanisms and regional and international spread of emerging MDR and XDR strains[26]. Blood culture confirmation and isolate characterisation is a pre-requisite for such activities, but neither are routinely performed in laboratories in areas where typhoid fever is endemic. However, in many developed nations, *S*. Typhi infections are notifiable and the disease is primarily associated with travellers returning from high-risk regions[27], providing an opportunity for informal sentinel surveillance of those regions. In England, typhoid fever is notifiable and all *S*. Typhi isolates are sent to Public Health England (PHE) for confirmation and characterisation via WGS, and recent travel history is sought via enhanced surveillance questionnaires. Most cases identified in England (~80%) are associated with recent travel to typhoid endemic areas[27]. We recently reported that the geographic origin of travel-associated *S*. Typhi cases in London could be predicted by comparing WGS data to the global framework[11], and identified the first case of ESBL *S*. Typhi in England from a patient with travel to Pakistan[28], supporting the notion that WGS data on travel-associated cases has sentinel surveillance value. Here we analysed WGS data from *S*. Typhi isolates referred to PHE from three years of national surveillance between April 2014 and April 2017[29], and explored the distribution of lineages and AMR determinants in geographic regions frequented by UK travellers.

## Methods

### Bacterial isolates included in this study

A total of 533 *S*. Typhi isolates from English cases received by PHE during the period of 1^st^ April 2014 to 31^st^ March 2017 were included in this study. Patient travel information was available for 488/533 of the isolates and was obtained by PHE using an enhanced surveillance questionnaire (https://www.gov.uk/government/publications/typhoid-and-paratyphoid-enhanced-surveillance-questionnaire), this included questions pertaining to the destination of any foreign travel that occurred during the likely incubation period (28 days before onset of symptoms). For the remaining 45 isolates, no enhanced surveillance questionnaire was completed or the data collected was incomplete; these were spread across the four years (n = 15 for 2014, n = 24 for 2015, n = 4 for 2016 and n = 2 for 2017; mean 8.5%). Details of all 533 isolates are provided in **S1 Table**.

### Whole genome sequencing

WGS was conducted as part of routine sequencing of all *Salmonella* isolates referred to the Gastrointestinal Bacteria Reference Unit at PHE, as previously described[29]. Briefly, DNA was fragmented and tagged for multiplexing with NexteraXT DNA Sample Preparation Kits (Illumina) and sequenced at PHE on a HiSeq 2500 yielding 100 bp paired end reads. FASTQ data is available from the NCBI Short Read Archive, BioProject accession PRJNA248792. Individual accession numbers for isolates analysed in this study are given in **S1 Table**.

### Single nucleotide variant (SNV) analysis and *in silico* genotyping

Paired end Illumina reads were mapped to the CT18 reference genome (accession AL513382)[30], which is the standard reference for *S*. Typhi genomic studies, using RedDog (V1.beta10.3) available at (https://github.com/katholt/RedDog). Briefly, RedDog maps reads to the reference genome using Bowtie2 (v.2.2.9)[31], before using SAMtools (v1.3.1)[32] to identify high quality single nucleotide variant (SNV) calls as previously described[19]. A core SNV alignment was generated for all SNV loci with consensus base calls (phred score >20) in >95% of genomes; this alignment was filtered to exclude SNVs in phage regions and repetitive sequences (354 kb; ~7.4% of bases in the CT18 reference chromosome as defined previously[10]; **S2 Table**) and recombinant regions identified by Gubbins[33]. The resulting alignment of 8053 SNVs was used as input to RAxML (v8.2.8) to infer a maximum likelihood (ML) phylogeny with a generalised time-reversible model and a Gamma distribution to model site-specific rate variation (GTR+ Γ substitution model; GTRGAMMA in RAxML), and 100 bootstrap pseudo-replicates to assess branch support.

Genotypes were inferred for all isolates by screening the Bowtie2 alignment (bam) files for SNVs used in the extended *S*. Typhi typing framework, GenoTyphi (code available at https://github.com/katholt/genotyphi)[11]. GenoTyphi uses 72 specific SNVs to assign isolates to one of four primary clusters; 16 clades; and 49 subclades[11], with the globally disseminated 4.3.1 (H58) subclade further delineated into lineages I and II (4.3.1.1 and 4.3.1.2).

### *In silico* characterisation of AMR associated genes and mobile elements

*S*. Typhi genomes were screened for acquired AMR genes and plasmid replicons using PHE’s Genefinder pipeline as previously described[34]. Reference sequences in Genefinder were curated from those described in Comprehensive Antimicrobial Resistance Database[35] and PlasmidFinder[36]. Genes were called as present within a genome when detected with 100% coverage and >90% nucleotide identity to the reference gene. Genefinder was also used to detect point mutations in the QRDR of chromosomal genes *gyrA* (codons 83, 87) and *parC* (codons 79, 80, 84)[37,38]. Isolates were defined as being MDR if genes were detected by Genefinder in the Beta-Lactamases, Trimethoprim, Sulphonamides and Chloramphenicol classes.

ISmapper[39] (v2) was run with default parameters to screen all read sets for insertion sites of the transposase IS*1* (accession X52534) relative to the CT18 reference chromosome sequence (accession AL513382). The binary data from ISmapper was processed in R using tidyverse (v1.2.1) (https://CRAN.R-project.org/package=tidyverse). For six isolates that carried MDR genes but no plasmid replicon genes, ISMapper was re-run with the minimum read depth threshold reduced to 1 read (from the default of 6 reads), in order to increase sensitivity to detect the associated IS*1* site. This identified IS*1* sites in five of the six isolates (SRR3049053, SRR5989319, SRR1967049, SRR5500465 and SRR5500455) but not in isolate SRR5500435.

The presence and subtypes of IncHI1 and IncN plasmids was further investigated using plasmid MLST (pMLST). Publicly available pMLST schemes for IncHI1[40] and IncN[41], available at https://pubmlst.org/plasmid/, were used to screen the relevant read sets using SRST2 (v0.1.8)[42]. The five isolates that were determined to have pST2 versions of the IncHI1 plasmid were then mapped to the reference genome of plasmid pAKU1 (accession AM412236) with RedDog along with a pST1 plasmid sequence (pUI1203_01 accession ERR340785) from *S*. Typhi strain UI1203 (genotype 3.2.1, isolated from Laos in 2001)[11] as an outgroup for phylogenetic tree rooting. The resulting SNV alignment was filtered to exclude non-backbone regions of the plasmid (66,458 bp), sites present in <95% of plasmid sequences, and recombinant regions detected by Gubbins (v2.3.2), resulting in a final alignment of 17 SNVs for phylogenetic inference as above.

Genomes with less common AMR profiles were assembled using Unicycler v0.4.6 [43]. The assemblies were further interrogated for the presence of plasmid replicon genes using ABRicate (https://github.com/tseemann/abricate) and the PlasmidFinder database using a minimum coverage and minimum identity of 90. The location of AMR genes detected, IS*1* sequences and IncHI1 plasmid backbone genes was visualised in Bandage v0.8.1 [44].

Trees annotated with MDR, IS*1* insertion sites, plasmid replicons and QRDR point mutations were visualised using ggtree v1.8.1[45]. An interactive version of the *S*. Typhi ML phylogeny with associated metadata is found at microreact (https://microreact.org/proiect/r1niJ_qJ4).

### Ciprofloxacin susceptibility phenotyping

The minimum inhibitory concentration (MIC) of ciprofloxacin was measured for 173 isolates (n = 117 in 2015 and n = 56 from 2016) as part of a previously reported study[37]. Briefly, MIC was determined by agar dilution using Mueller-Hinton agar, and *S*. Typhi were interpreted as displaying resistance (MIC ≥0.5 μg/mL) or susceptibility to ciprofloxacin (MIC <0.06 μg/mL), according to the EUCAST guidelines (v7.1)[37]. For the 173 phenotyped isolates, the total number of QRDR mutations per isolate was plotted against the reported MIC, in R using ggplot2 (v3.0.0)[46].

## Results

### Countries of travel

A total of 533 *S*. Typhi isolates were referred from English cases to PHE during the three-year period between April 2014 and April 2017 (n=178, n=176, n=179 in each year running from April to March; listed in **S1 Table**). Of these, 449 cases (84.2%) reported recent foreign travel (within 28 days of onset of symptoms), spanning 26 countries. A further 39 cases (7.3%) reported no travel abroad. The proportion of cases reporting no recent travel, which may represent local transmission within the UK, displayed a non-significant increase from 5.6% in the first year to 8.9% in the third year (p=0.5 using Chi-squared trend test). No information on travel was available for the remaining 45 cases (8.4%). The distribution of travel destinations for the cases with known travel history are given in **Table 1** and **Fig. 1**. The majority of cases (n=387, 73%) reported travel to South Asia, particularly India (36%), Pakistan (29%) and Bangladesh (6%), reflecting travel and migration patterns between England and typhoid endemic areas. Annual case numbers from India and Bangladesh were constant (mean 64 and 11 per year, respectively; p=0.2 in each case using Chi-squared trend test), but case numbers from Pakistan spiked significantly in the third year of the study, to n=68 (38% of total cases) compared to n=45 and n=43 (25%) in the first two years (p=0.01 using Chi-squared trend test). Small numbers of cases reported travel to South East Asia (n=10, 2%), East Africa (n=21, 4%), West Africa (n=11, 2%) and the Middle East (n=4, 1%; see **Table 1**). Five cases (1%) had reported travel to Europe or North America or South America only.

**Table 1:** Summary table of *S*. Typhi collection.

|  | Total | 4.3.1 (%) | MDR (%) | Number of QRDR mutations |  |  |
| --- | --- | --- | --- | --- | --- | --- |
| Country |  |  |  | None (%) | One or two (%) | Three (%) |
| South Asia |  |  |  |  |  |  |
| Afghanistan | 1 | 1 (100%) | 1 (100%) | 0 (0%) | 1 (100%) | 0 (0%) |
| Bangladesh | 33 | 17 (51.5%) | 8 (24.2%) | 2 (6.1%) | 31 (93.9%) | 0 (0%) |
| India | 191 | 151 (79.1%) | 6 (3.1%) | 5 (2.6%) | 143 (74.9%) | 43 (22.5%) |
| Nepal | 3 | 1 (33.3%) | 0 (0%) | 1 (33.3%) | 1 (33.3%) | 1 (33.3%) |
| Pakistan | 156 | 143 (91.7%) | 85 (54.4%) | 10 (6.4%) | 140 (89.7%) | 6 (3.9%) |
| South East Asia |  |  |  |  |  |  |
| Indonesia | 3 | 0 (0%) | 0 (0%) | 3 (100%) | 0 (0%) | 0 (0%) |
| Malaysia | 1 | 1 (100%) | 0 (0%) | 0 (0%) | 1 (100%) | 0 (0%) |
| Myanmar | 2 | 1 (50%) | 0 (0%) | 0 (0%) | 2 (100%) | 0 (0%) |
| Philippines | 4 | 0 (0%) | 0 (0%) | 4 (100%) | 0 (0%) | 0 (0%) |
| **Middle East** |  |  |  |  |  |  |
| Iraq | 3 | 1 (33.3%) | 0 (0%) | 2 (66.7%) | 1 (33.3%) | 0 (0%) |
| UAE | 1 | 1 (100%) | 1 (100%) | 0 (0%) | 1 (100%) | 0 (0%) |
| **East Africa** |  |  |  |  |  |  |
| Ethiopia | 1 | 0 (0%) | 0 (0%) | 1 (100%) | 0 (0%) | 0 (0%) |
| Mozambique | 1 | 1 (100%) | 1 (100%) | 1 (100%) | 0 (0%) | 0 (0%) |
| Rwanda | 1 | 1 (100%) | 0 (0%) | 0 (0%) | 1 (100%) | 0 (0%) |
| Tanzania | 4 | 4 (100%) | 2 (50%) | 2 (50%) | 2 (50%) | 0 (0%) |
| Uganda | 4 | 4 (100%) | 0 (0%) | 0 (0%) | 4 (100%) | 0 (0%) |
| Zimbabwe | 10 | 8 (80%) | 8 (80%) | 8 (80%) | 2 (20%) | 8 (80%) |
| **West Africa** |  |  |  |  |  |  |
| Angola | 2 | 0 (0%) | 0 (0%) | 2 (100%) | 0 (0%) | 0 (0%) |
| Ghana | 2 | 0 (0%) | 1 (50%) | 2 (100%) | 0 (0%) | 0 (0%) |
| Nigeria | 6 | 0 (0%) | 2 (33.3%) | 3 (50%) | 3 (50%) | 0 (0%) |
| Sierra Leone | 1 | 0 (0%) | 0 (0%) | 1 (100%) | 0 (0%) | 0 (0%) |
| **Other Regions** |  |  |  |  |  |  |
| China | 1 | 0 (0%) | 0 (0%) | 0 (0%) | 1 (100%) | 0 (0%) |
| Egypt | 1 | 0 (0%) | 0 (0%) | 0 (0%) | 1 (100%) | 0 (0%) |
| Greece | 1 | 0 (0%) | 0 (0%) | 0 (0%) | 1 (100%) | 0 (0%) |
| Peru | 1 | 0 (0%) | 0 (0%) | 0 (0%) | 1 (100%) | 0 (0%) |
| USA | 1 | 0 (0%) | 0 (0%) | 1 (100%) | 0 (0%) | 0 (0%) |
| **No Travel** | 39 | 15 (38.5%) | 5 (12.8%) | 19 (48.7%) | 18 (46.2%) | 2 (5.1%) |
| **Unknown** | 45 | 30 (66.7%) | 9 (20%) | 9 (20%) | 32 (71.1%) | 4 (8.9%) |

UAE: United Arab Emirates
MDR: Multi-drug resistance
QRDR: Quinolone resistance determining region

**Fig. 1.**
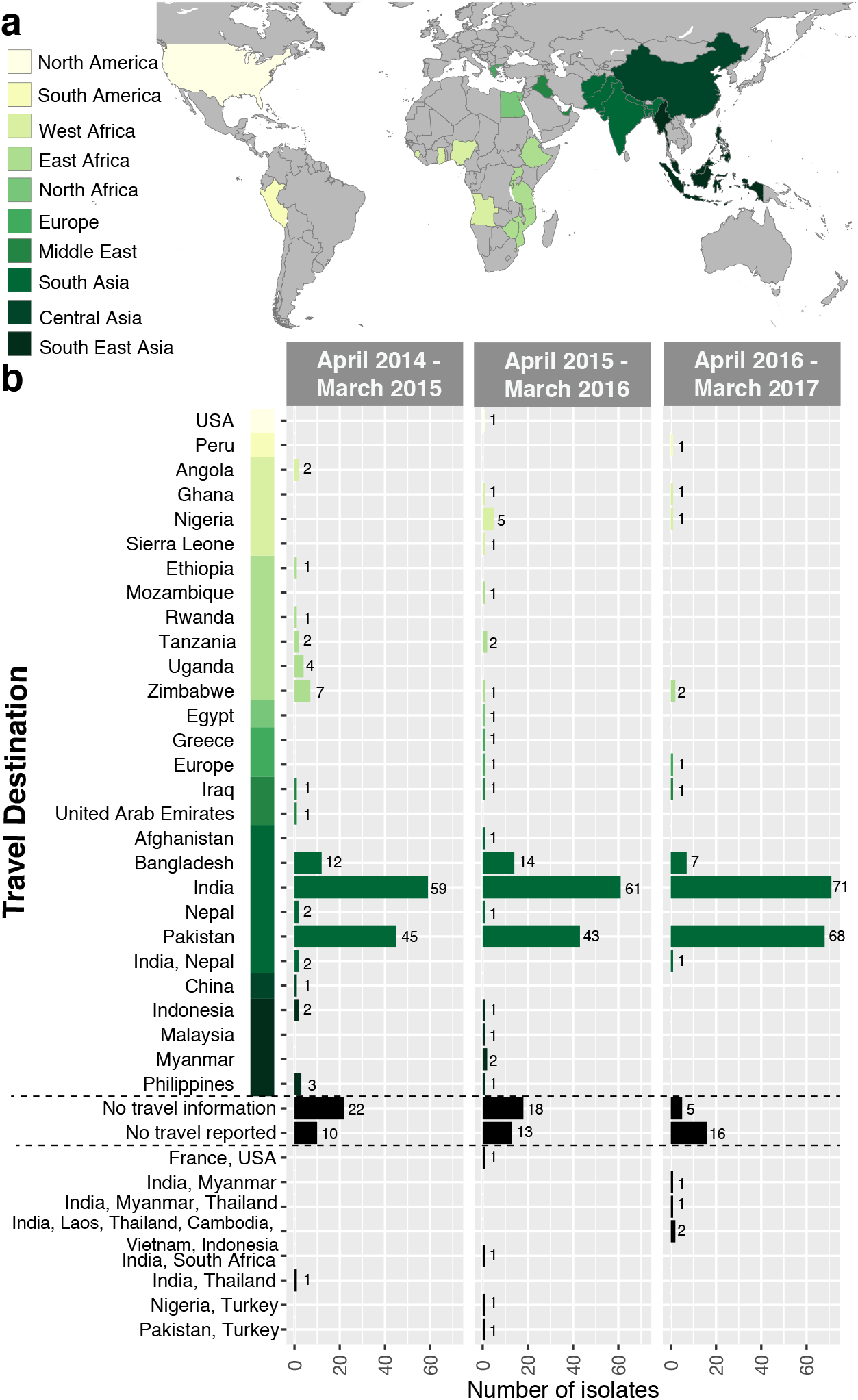
Summary of *S*. Typhi received at Public Health between April 2014 to March 2017. a) Map highlighting countries that were reported as travel destinations for patients from which *S*. Typhi cultures were collected. Countries are coloured by major geographical regions. b) Number of *S*. Typhi isolates reporting travel to each country over the three-year period of study, stratified by date of receipt at PHE.

### *S*. Typhi genotypes by region

The 533 genomes were assigned to 31 unique *S*. Typhi genotypes using the GenoTyphi scheme (**Fig. 2**). The majority (n=391, 73.4%) belonged to subclade 4.3.1 (H58), which was identified in cases with travel to 15 different countries. Genotype 4.3.1 dominated amongst cases with reported travel to the South Asian countries associated with the greatest burden of travel-related cases (India, 79% 4.3.1; Pakistan, 92% 4.3.1; Bangladesh, 52% 4.3.1), and amongst cases with travel to East Africa (86% 4.3.1; see **Table 1**). Of the 4.3.1 isolates, 162 (41%) were further classified to lineage I (4.3.1.1), including 66% of the 4.3.1 isolates from cases with travel to Pakistan, 94% of those with travel to Bangladesh, and all 4.3.1 isolates from cases with travel to Mozambique, Tanzania and Zimbabwe (total n=13). In contrast, lineage II (4.3.1.2) accounted for 156 (40%) of 4.3.1 genotypes, dominating in returning travellers to India (75% of 4.3.1 isolates) and accounting for the three 4.3.1 isolates associated with travel to Rwanda and Uganda (**Fig. 3**). Notably, 4.3.1 was not detected in cases associated with travel to West Africa (0/11; see **Table 1, Fig. 3**). Other common genotypes include clade 3.3 (6% of total isolates), found in cases with travel to India (9% of all isolates from this location), or Bangladesh (15%); clade 2.2 (3.1% of total isolates), found in cases with travel to India (2%) or Pakistan (4.5%); subclade 3.2.2 (2.6% of total isolates), found in 21% of cases with travel to Bangladesh; and subclade 3.1.1 (1.7% of total isolates), found in 7/11 (63%) of cases with travel to West Africa, including Nigeria (n=4/6), Ghana (2/2) and Sierra Leone (1/1).

**Fig. 2.**
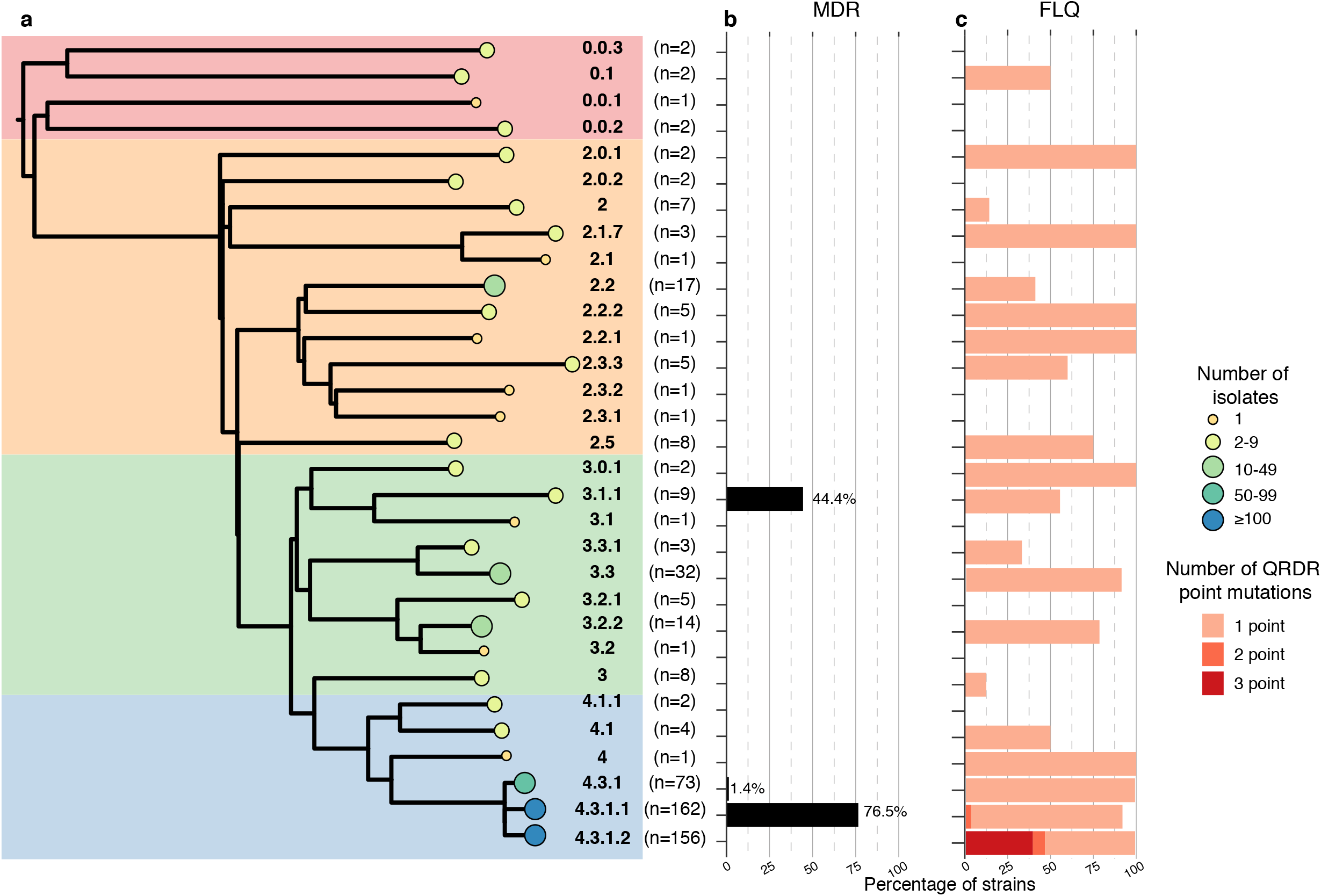
Membership of *S*. Typhi by genotypes and association with AMR. a) Population framework tree of the *S*. Typhi genotypes sequenced at PHE. Background shading indicates the primary clades. Tree tips indicate individual genotypes assigned by GenoTyphi; tip sizes and colours indicate number of isolates belonging to each genotype, these values (n) are also printed to the right. b) Frequency of MDR for each genotype (defined based on presence of AMR genes encoding resistance to the first line drugs ampicillin, trimethoprim-sulfamethoxazole, or chloramphenicol). c) Frequency of QRDR mutations by genotype.

**Fig. 3.**
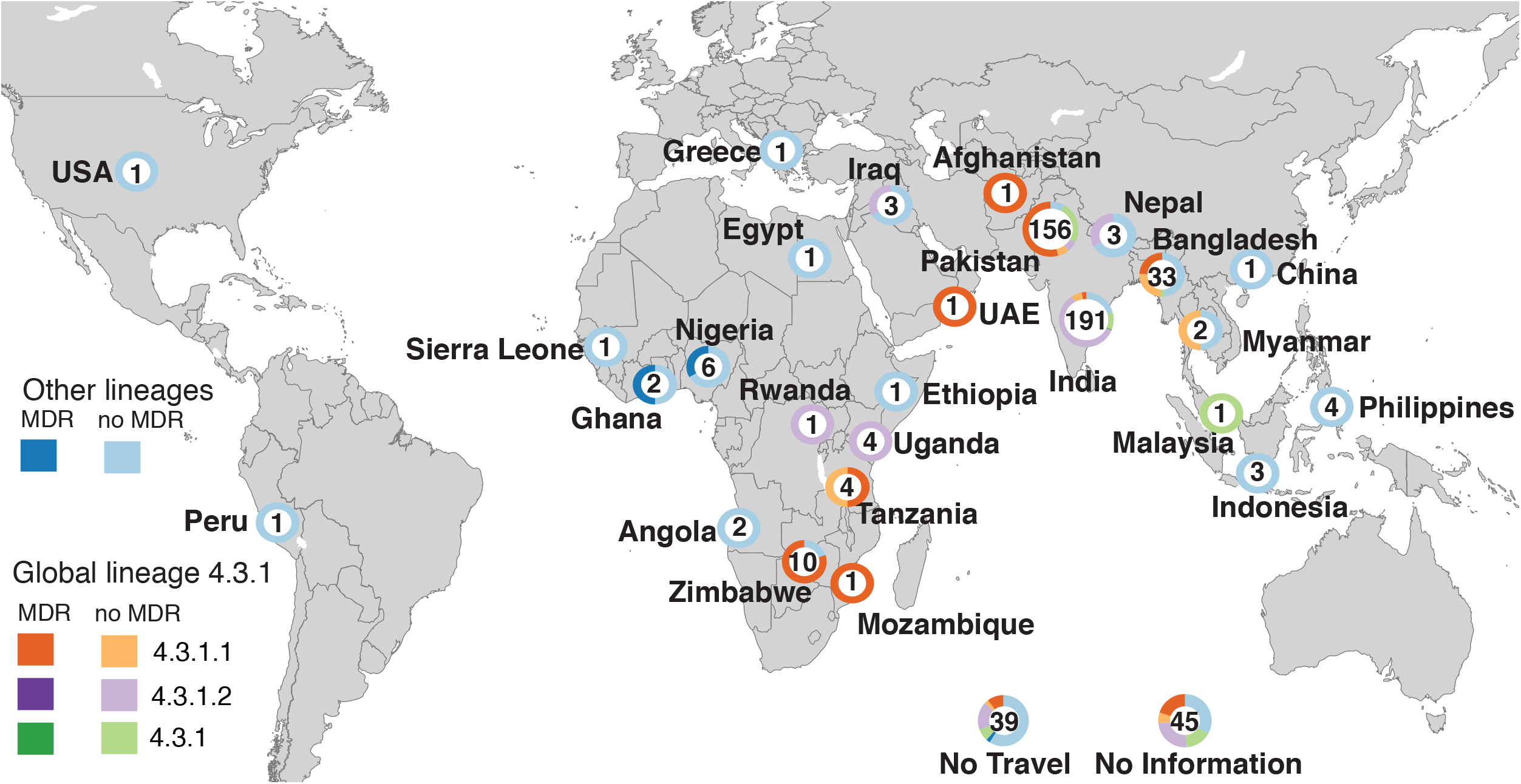
Geographic distribution of *S*. Typhi based on reported country of travel. *S*. Typhi isolates with completed travel questionnaires and reporting travel to only one country are shown. The number in each circle indicates the total number of cases reporting travel to that country, circle graphs indicate the genotype composition of the corresponding *S*. Typhi isolates, stratified by MDR, coloured as per inset legend.

### Multi-drug resistance and QRDR mutations by region

The overall frequency of MDR, defined as carriage of genes associated with resistance to ampicillin (*bla_TEM-1_*), chloramphenicol (*catA1*) and co-trimoxazole (*sul1* or *sul2* plus a *dfrA* gene), was 129 (24%). MDR *S*. Typhi was associated with reported travel to 10 countries, with high frequencies amongst isolates whose cases report travel to Zimbabwe (80%), Nigeria (33%), Tanzania (50%), Ghana (50%), Bangladesh (24%) and Pakistan (55%) (**Table 1, Fig. 3**). MDR was also present at low frequency amongst cases with reported travel to India (3%), and singleton MDR isolates were detected from Mozambique, United Arab Emirates and Afghanistan. Notably the majority of MDR isolates were linked with travel to Pakistan (n=85, 66% of all MDR isolates) and other South Asian countries (n=100, 77% of all MDR isolates; see **Table 1**).

All MDR isolates belonged to either 4.3.1 (n=125) or 3.1.1 (n=4, see **Fig. 2b**). The frequency of MDR was highest in 4.3.1.1 (96%), then 3.1.1 (3%), and one isolate in 4.3.1 (**Fig. 2b, Fig. 3**). At the regional level, MDR was common amongst cases associated with travel to East Africa (52%), West Africa (27%) and South Asia (26%), but not with travel elsewhere (with the exception of one isolate from UAE; see **Table 1**). However, there were significant differences between countries within these regions, associated with differences in the dominant *S*. Typhi clades (**Fig. 3**). In East Africa and South Asia, MDR was detected amongst cases associated with travel to the countries dominated by MDR-associated lineage 4.3.1 lineage I (Mozambique, Tanzania and Zimbabwe in East Africa; and Bangladesh and Pakistan in South Asia; see **Table 1, Fig. 3**). In West Africa, MDR was detected in three isolates (n=2 Nigeria, n=1 Ghana), all belonging to the region’s dominant subclade 3.1.1 (a fourth MDR 3.1.1 infection had no reported travel).

QRDR mutations were identified in 455 isolates (85.4%) originating from 18 countries (**Table 1, S1 Table**). QRDR mutants were most frequent in India (97%), Bangladesh (94%), Nepal (66%), Pakistan (94%), Myanmar (100%), Uganda (100%), and Nigeria (50%). Singleton QRDR mutants were also identified in China, Malaysia, Afghanistan, United Arab Emirates, Egypt, Rwanda, Peru, Greece and Zimbabwe. QRDR triple mutants were identified only in isolates associated with travel to South Asia: 33% of isolates from Nepal, 23% of those from India and 4% of those from Pakistan (**Table 1**). Overall, the presence of QRDR mutations was very common in cases associated with travel to South Asia (95% of all isolates linked to this region), and significantly less common (p < 2.2e16 using Chi-squared trend test) in East Africa (43%,) and West Africa (27%; see **Table 1**).

Ciprofloxacin MICs have been previously reported for 173 of the *S*. Typhi isolates (**S1 Table**)[37], and a comparison of QRDR mutations with these phenotypes is shown in **Fig. 4** to facilitate the interpretation of QRDR mutations. These data showed that all isolates carrying a single QRDR mutation had ciprofloxacin MIC of at least 0.064 μg/mL (exceeding the EUCAST threshold for susceptibility); all those with two QRDR mutations had MIC of 0. 125 μg/mL; and all those with three mutations had MIC ≥1 μg/mL (above the threshold for resistance) (**Fig. 4**). The most common mutations were *gyrA*-S83F (n=271, 51%) and *gyrA-* S83Y (n=83, 16%), which were found in 18 genotypes in cases associated with travel to eleven and six countries respectively (**Table 2**). Double mutants (*gyrA-S38F* or -S83Y plus a mutation in *parC*), associated with elevation of ciprofloxacin MIC to ≥0.25 μg/mL (**Fig. 4**), were found in 16 isolates (3.0%) associated with travel to India or Bangladesh. A total of 61 isolates were identified as QRDR triple mutants, which display resistance to ciprofloxacin (MIC >1 μg/mL, see **Fig. 4**) and have been associated with fluoroquinolone treatment failure[8]. Most of these (n=59) carried gyrA-S83F, *gyrA* -D87N and *parC-S80I* mutations, while the remaining two had a unique profile of *gyrA-S83F, gyrA-D87G* and *parC-S80I* and of *gyrA-S83FY, parC-Y74X* and *parC-P98X* (**Table 2**).

**Table 2:**
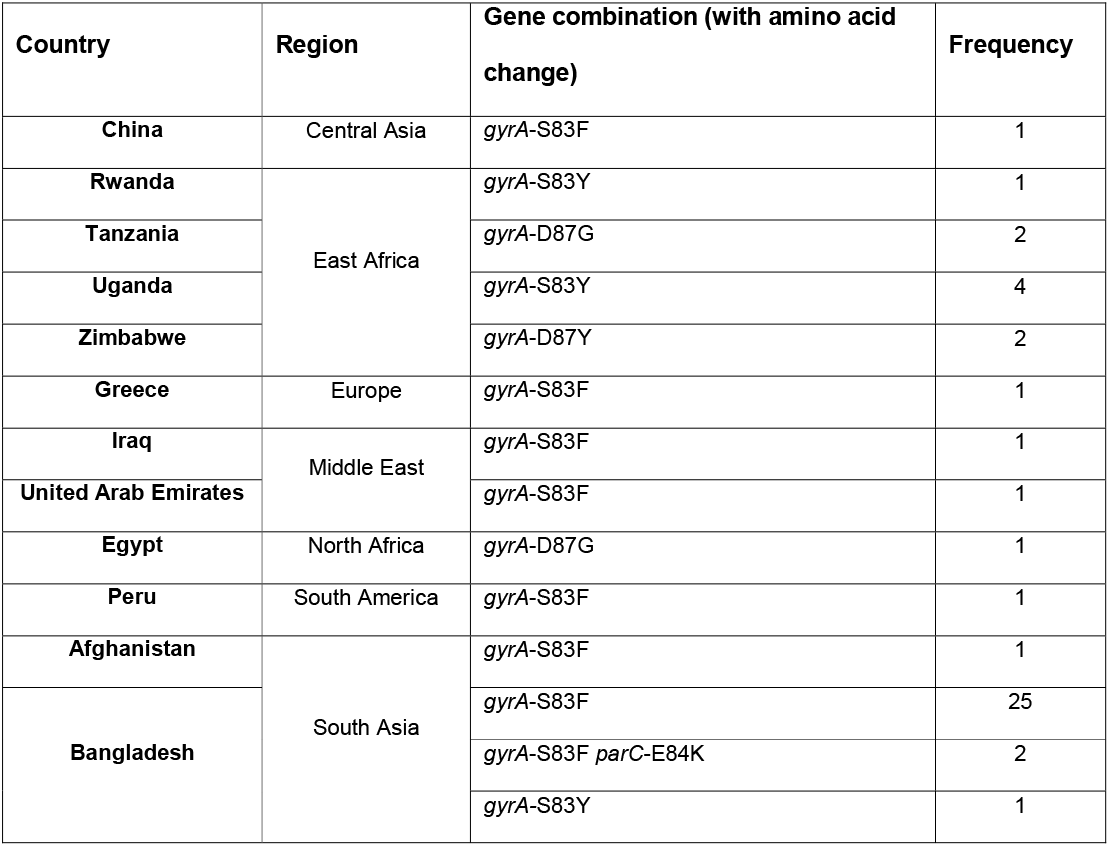
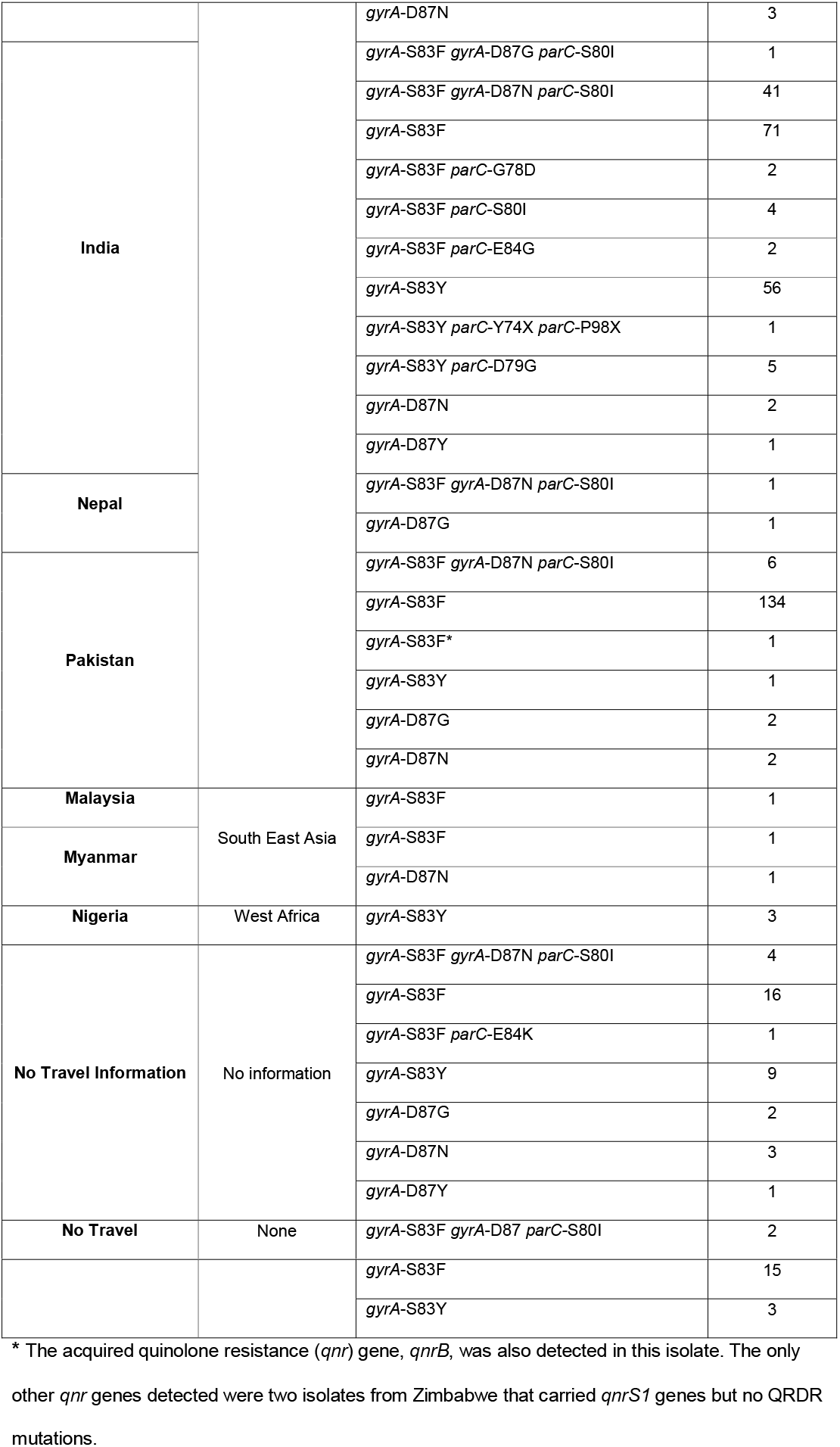
Combinations of coding changes detected in the QRDR and their frequency by country of reported travel.

**Fig. 4.**
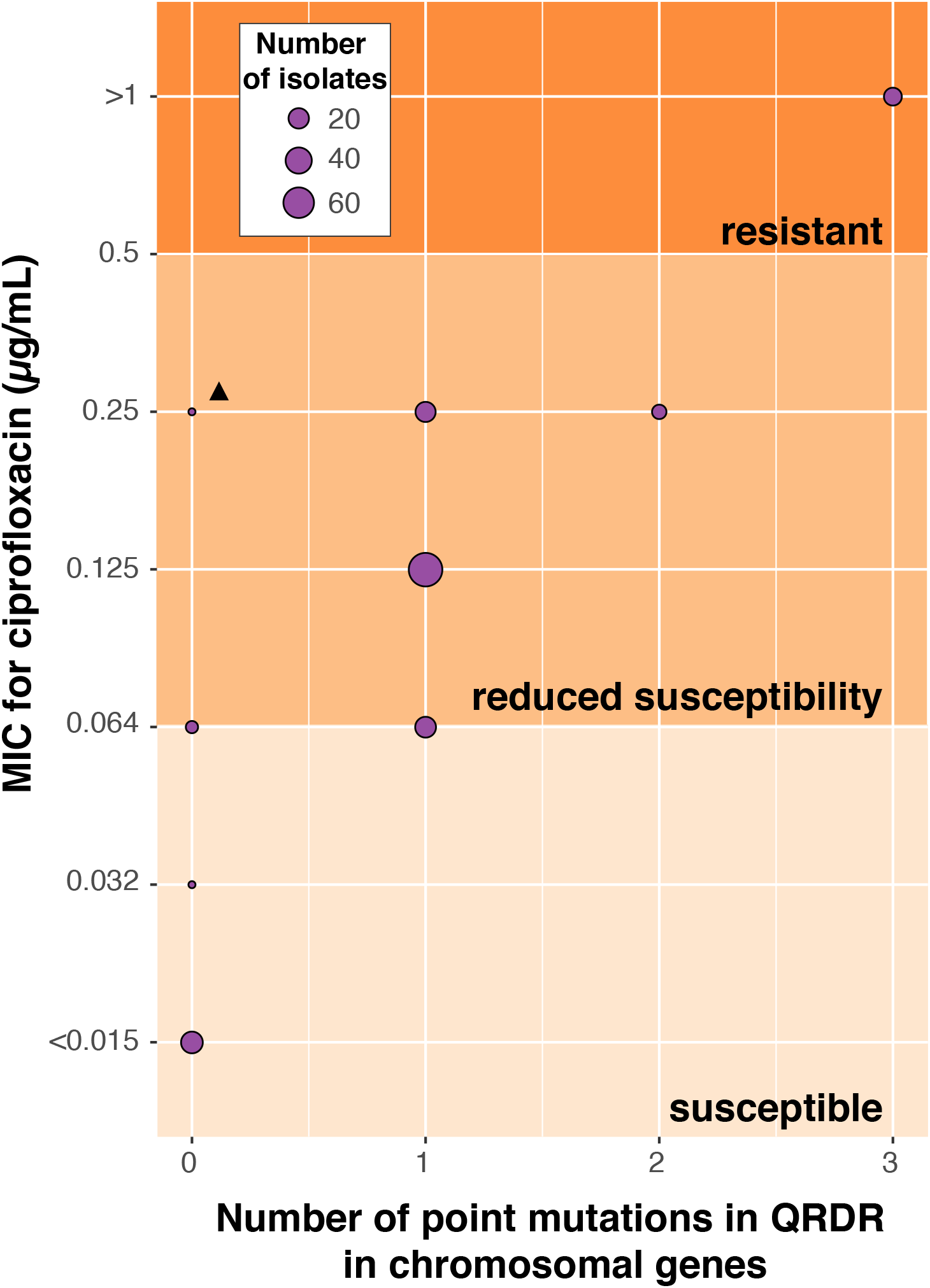
Minimum inhibitory concentration (MIC) for ciprofloxacin versus number of point mutations in QRDRs in two chromosomal genes. The number of isolates with different combinations of MIC values and point mutations detected in the QRDRs of two chromosomal genes, *gyrA* and *parC,* are shown. The breakpoints of susceptible, reduced susceptibility and resistance are shown by the background gradient of the plot. The triangle indicates an isolate that carries *qnrS-1*.

All QRDR double and triple mutants belonged to genotype 4.3.1 and were associated with travel to South Asia (or no/unknown travel). Overall, QRDR mutations were significantly more common in 4.3.1 lineage II compared to lineage I (99% vs 91%, p=0.006, two-sided test of difference in proportions). The 61 QRDR triple mutants were detected only in *S*. Typhi 4.3.1 lineage II isolates (Fig. 2c) and were associated with either travel to India (n=43, 22.5% of all isolates from this location, where lineage II is dominant); travel to neighbouring countries Pakistan (n=6) or Nepal (n=1); travel to multiple destinations including India (n=5); no reported travel (n=2); or no travel information available (n=4) (**Tables 1-2**). Notably because all 4.3.1 MDR isolates belonged to lineage I, and all QRDR triple mutants belonged to lineage II (**Fig. 2**), there were no isolates with both MDR and three QRDR mutations (**Fig. 2, S1 Table**). There were however 119 cases with MDR plus 1-2 QRDR mutations (i.e. reduced susceptibility to fluoroquinolones, **Fig. 4**). The vast majority of these were 4.3.1 lineage I (n=115), most commonly associated with travel to Pakistan (n=85, 54.4% of isolates from this country), Bangladesh (n=8, 24%) and India (n=6, 3%) but also occasional cases who reported travel to Zimbabwe (n=2), Tanzania (n=2) and United Arab Emirates (n=1). Three 3.1.1 isolates were also MDR with one QRDR mutation (n=2 travel to Nigeria, n=1 with no recorded travel).

### Plasmid vs chromosomal location of AMR genes in *S*. Typhi

All MDR isolates belonged to 4.3.1 (n=125) or 3.1.1 (n=4) and carried the typical *S*. Typhi MDR composite transposon comprising Tn6029 (encoding *bla_TEM-1_, sul2, strAB*) inserted in Tn*21* (carrying a class I integron encoding *dfrA* alleles in the gene cassette and *sul1* at the end), which is in turn inserted within Tn*9* (encoding *catA1*)[47] (see **Fig. 5a**). All 125 MDR 4.3.1 isolates (associated with South Asia and East Africa) carried *dfrA7* in the integron cassette and no plasmid replicons (**Fig. 5a**). In most of these (n=123, 98%), we detected chromosomal IS*1* insertions at sites previously associated with IS*1*-mediated integration of the MDR composite transposon (**S1 Fig.**) (*cyaA* or *yidA* sites[12,18]). A putative IS*1* insertion was detected in the novel site STY3168 in a single genome (SRR5500440). Further, of the six isolates with four IS*1* sites detected, five had recent travel to Pakistan with last isolate having no reported travel (**S1 Fig.**). Notably, most (93%) of the MDR 4.3.1 isolates also carried a QRDR mutation, the most common being *gyrA-S83F* (**Table 2, S1 Fig**). The four MDR 3.1.1 isolates (associated with West Africa) carried IncHI1 PST2 plasmids with *dfrA15* in the integron cassette (**Fig. 5b**). An IncHI1 PST2 plasmid was also identified in a single non-MDR 2.3.1 isolate associated with travel to Nigeria. The plasmid backbone was very closely related to that of the 3.1.1 West African plasmids but carried *dfrA7* in the integron cassette and lacked the chloramphenicol and ampicillin resistance genes *catA1* and *bla_TEM_* (**Fig. 5a-b**).

**Fig. 5.**
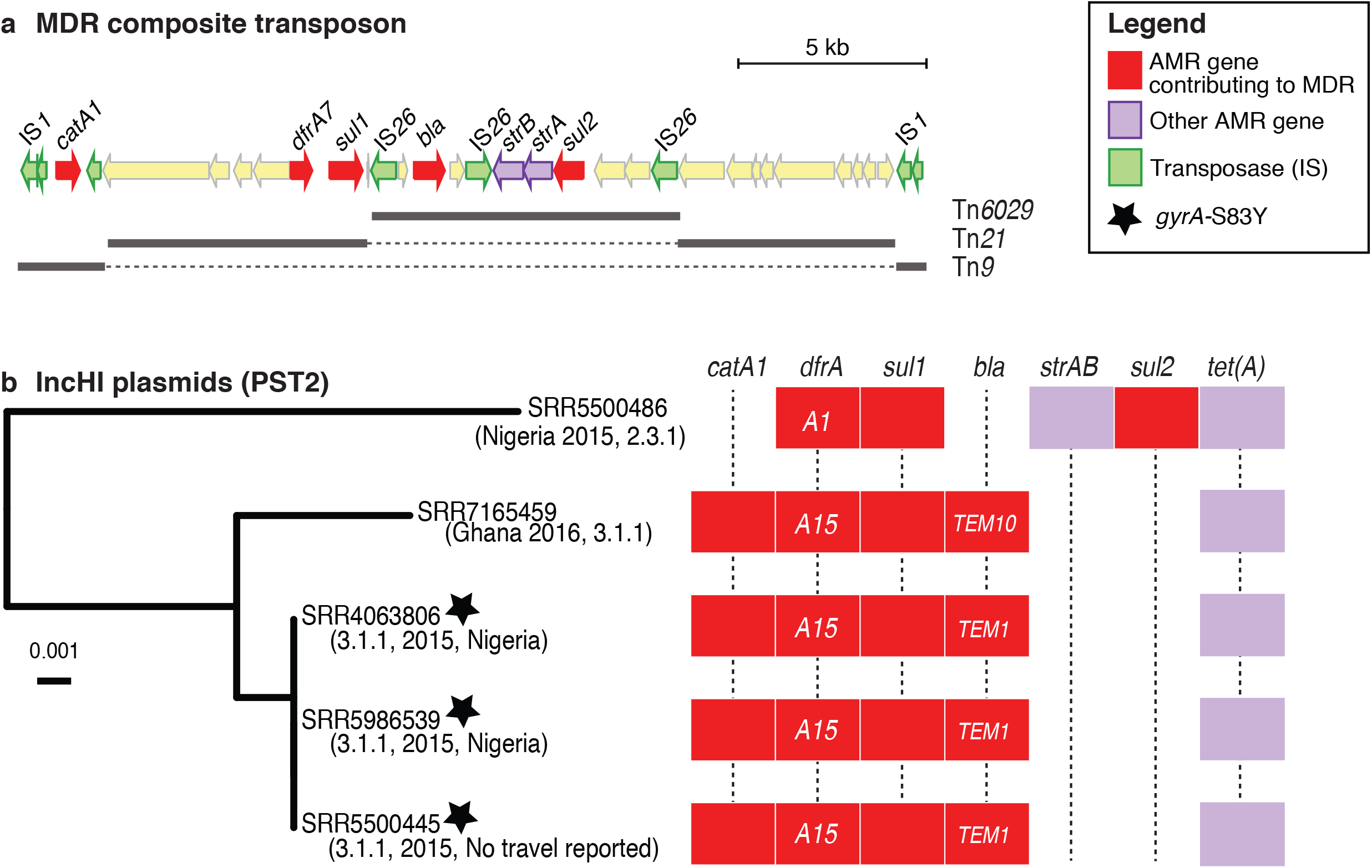
Mobilisation of MDR element in *S*. Typhi. a) Structure of the MDR composite transposon in *S*. Typhi strain ERL12960 (4.3.1 lineage I) is shown (accession ERL12960). MDR genes are those encoding resistance to the first line drugs ampicillin, trimethoprim-sulfamethoxazole or chloramphenicol b) Phylogeny showing genetic relationships between the five IncHI1 PST2 *S*. Typhi isolates identified in this study, based on SNVs identified in the plasmid backbone sequence and rooted using the PST1 plasmid pUI1203_01 (accession ERR340785) as an outgroup. The presence of AMR genes detected in each genome is indicated in the heatmap, with alleles specified for *dfrA* and *bla_TEM_* genes.

Plasmid replicon and AMR gene screening identified additional plasmid replicons and AMR genes at low frequency. Two of the MDR 4.3.1.1 isolates from Zimbabwe carried IncN (subtype PST3) plasmids. In addition to the genes typical of the MDR composite transposon (with *dfrA7* in the integron), these isolates also carried *qnrS, dfrA14* and *tet(A).* It was not possible to resolve the precise locations of the acquired AMR genes due to the limitations of short read assembly. As these isolates lack QRDR mutations, the presence of the *qnrS* gene is predicted to confer reduced susceptibility to fluoroquinolones but not full resistance; indeed, one of the isolates (SRR4063811) was phenotyped and displayed reduced susceptibility to ciprofloxacin, with MIC 0.25 μg/mL. IncN plasmids (subtype PST5) were found in three non-MDR 4.3.1.2 isolates that carried *dfrA15, sul1* and *tet(A).* These isolates (two from India, one with no reported travel) were also QRDR triple mutants and thus predicted to be fully resistant to fluoroquinolones. The combination of *dfrA14, sul2, bla_TEM-1_, strA* and *strB* was detected in three 4.3.1.1 isolates with no QRDR mutations (two with travel to Tanzania, one with no travel information). All three carried sequences with similarity (100% nucleotide identity and 50% coverage) to a FIB_K_ plasmid carrying *dfrA14, sul2* and *bla_TEM-1_* that was previously sequenced from a 2008 Tanzanian *S*. Typhi, strain 129-0238 (GenBank accession LT904889)[12]. The same combination of AMR genes (*dfrA14, sul2, bla_TEM-1_, strA* and *strB*) plus *tet(A)* were also found in a 3.1.1 *gyrA-S83Y* isolate from Nigeria that harboured an IncY plasmid replicon.

## Discussion

Here we demonstrate the utility of using *S*. Typhi WGS data generated routinely at a public health laboratory in the UK to serve as informal surveillance for different geographical regions where typhoid fever is endemic. The PHE dataset encompasses a diverse collection of 533 *S*. Typhi isolates from multiple geographical regions collected over a three-year sampling period (**Fig. 1**). As typhoid fever has not been endemic in England since the successful interventions of the major controlling measures, water sanitation and hygiene complemented with antimicrobial therapy[27,48], it is assumed that notified cases to PHE are associated with returned travellers and their contacts. The collection is biased towards isolates from South Asia and East Africa which reflects historical and contemporary ties between England and these regions, however these are also regions that experience a high burden of typhoid fever and could benefit from the AMR and WGS data obtained routinely by PHE. Notably, public health agencies in other countries receive *S*. Typhi isolates reflecting the distinct travel habits of their own populations (for example Institut Pasteur receives more *S*. Typhi from travellers visiting Francophone countries in Africa and former French colonies such as Vietnam), and the synthesis of these diverse collections could potentially provide more extensive sentinel surveillance coverage of typhoid endemic regions.

Typing the *S*. Typhi isolates using the GenoTyphi scheme enabled rapid classification of the 533 genomes into 31 distinct lineages (**Fig. 2**). This simple tree-free approach showed clustering of subclades by geographical region of travel that was consistent with previous previously reported geographical patterns[11], providing support for the use of travel associated isolates as an indicator of local pathogen populations. Notable examples include the detection of MDR subclade 4.3.1 in East Africa and South Asia [8,15], and the complete absence of subclade 4.3.1 in the isolates from West Africa[14] (**Fig. 3**). Genomes from cases reporting travel to West Africa were genotyped as 3.1.1, consistent with earlier studies where 3.1.1 was found to be the main *S*. Typhi lineage in the region[13,14]. Furthermore, for isolates with travel to more than one country, the genotype can help to discern the most likely origin of the infection; for example, for one case reporting travel to Nigeria and Turkey, the genome isolate (SRR558502) was identified as genotype 3.1.1, suggesting that the pathogen was most likely acquired in Nigeria. This further demonstrates the public health utility of WGS data on returning traveller isolates. Notably the GenoTyphi scheme provides a mechanism and nomenclature for such insights to be achieved simply and rapidly from individual genomes and by different laboratories working independently, without need for phylogenetic tree construction or other comparative analyses.

Data from this study revealed that reduced susceptibility to fluoroquinolones was common amongst *S*. Typhi associated with diverse geographic sites and genotypes. However, the ciprofloxacin resistant QRDR triple mutants were all from cases belonging to subclade 4.3.1.2 (**Fig. 2, Table 2**), the majority of which had reported travel to South Asia, mainly India. These data align with previous findings[8,19], and support the hypothesis that high levels of fluoroquinolone exposure in India through healthcare and the environment are driving the emergence of resistant *S*. Typhi and other pathogens in the region[49] and contributing to treatment failure for typhoid fever[8]. Further, in addition to confirming that QRDR triple mutants are resistant to ciprofloxacin, the MIC data clearly showed that even a single QRDR mutation is associated with reduced ciprofloxacin susceptibility (MIC 0.06-0.25 μg/mL) (**Fig 4**), which has also been associated with clinical failure[37,50].

Strikingly in this data set all 4.3.1 MDR isolates carried the composite transposon integrated in the chromosome, suggesting an enormous shift in the burden of MDR typhoid from plasmid-borne resistance. This is of grave concern as it means that there is likely to be very little fitness cost associated with carriage of the MDR transposon. In the late 1990s the increase in MDR typhoid in Asia prompted a switch to fluoroquinolones for treatment, which in Nepal and other regions was followed by almost complete loss of the MDR plasmid from the *S*. Typhi population, suggesting that the fitness cost of the plasmid leads to plasmid loss in the absence of selection from the first-line drugs. However, the integration of the MDR transposon into the *S*. Typhi chromosome likely alleviates any fitness cost, making it more likely that MDR will be maintained even in the absence of selection for the specific resistances encoded. We showed that the two most common sites for chromosomal integration of the MDR element was at the known sites *yidA* or upstream of *cyaA* in the 4.3.1.1 lineage. Of particular note was the increase in number of IS*1* insertions in from cases in subclade 4.3.1.1 with reported travel to Pakistan (**S1 Fig.**). We may hypothesise that the *S*. Typhi from this region may be more likely to acquire novel mechanisms in response to local selective pressures. Indeed, the recent acquisition of an IncY plasmid harbouring *bla_CTX-M-15_* and *qnrS* genes has resulted in the emergence of an XDR lineage of *S*. Typhi from Pakistan[24] and an IncI1 plasmid encoding *bla_CTX-M-15_* in an *S*. Typhi from Bangladesh[51] provides evidence for this hypothesis, highlighting the importance of ongoing surveillance of these regions that experience a high burden of typhoid fever.

Currently there are three critical AMR threats posed by *S*. Typhi, namely the dissemination of mobile AMR genes mediating MDR profiles, the evolution of point mutations in *gyrA* and *parC,* two core housekeeping genes, that confer differing levels of fluoroquinolone resistance (**Fig. 2, Fig. 4**), and the recent emergence of XDR *S*. Typhi. The WGS data presented here provide insight into changing AMR dynamics within *S*. Typhi. Importantly, the concordance of genomic and phenotypic AMR data for ciprofloxacin resistance in this study (**Fig. 4**) which have been extensively characterised for *S*. Typhi for multiple drugs previously[37], demonstrates the utility of WGS for robustly characterising AMR profiles. Here, two of the AMR threats where characterised within the *S*. Typhi collection, reflecting geographical differences in AMR profiles. While no XDR *S*. Typhi had been detected in the PHE collection between April 2014 and March 2017, the first XDR *S*. Typhi isolate in a returned traveller with recent travel to Pakistan shortly after XDR *S*. Typhi were reported from Pakistan has been identified[24,28]. This highlights the value in using these data as informal surveillance of *S*. Typhi as we hypothesise that resistance to second-line drugs such as azithromycin will arise under continued selective pressure as has occurred with previous drugs.

## Funding information

DJI was supported by the Victoria Fellowship from the Department of Economic Development, Jobs, Transport and Resources, State Government of Victoria, Australia.

KEH was supported by a Senior Medical Research Fellowship from the Viertel Foundation of Australia and the Bill and Melinda Gates Foundation, Seattle. ZAD is supported by a project funded by the Wellcome Trust of Great Britain (106158/Z/14/Z). TJD is affiliated to the National Institute for Health Research Health Protection Research Unit (NIHR HPRU) in Gastrointestinal Infections at University of Liverpool in partnership with Public Health England (PHE), in collaboration with University of East Anglia, University of Oxford and the Quadram Institute.

## Conflict of interest

The authors declare no competing interests

## Author contributions

D.J.I., S.N., M.A.C., K.E.H. and T.J.D. contributed to the design of the study and data interpretation. D.J.I. performed the majority of bioinformatic analyses with input from Z.A.D., P.A., H.H., S.N, T.J.D. and K.E.H. The travel data was collected by J.F. The ciprofloxacin phenotypic resistance data was generated by S.N. and M.D. All authors contributed to the writing of the manuscript.

## Supporting information

**S1 Fig. Overview of *S*. Typhi PHE collection**

a) The phylogeny of the 533 *S*. Typhi isolates in the PHE collection is shown on the left with the geographical region of reported travel. b) The presence of IncHI1 or IncN plasmid replicons is shown with the different colours indicating PST. c) The presence of MDR profiles is shown in black. The presence of point mutations in QRDR genes is shown with the different gradients of blue indicating different mutations. d) Detected IS*1* sites of insertion are shown in black. The total number of IS detected in each of the *S*. Typhi isolates is shown on the far right.

**S1 Table. Data of 533 *S*. Typhi isolates received and sequenced at Public Health England between April 2014 and April 2017**

**S2 Table. Excluded repeat and phage regions in CT18 reference**

