## Supplementary figures and images for "Informal genomic surveillance of regional distribution of *Salmonella* Typhi genotypes and antimicrobial resistance via returning travellers"

### Supplementary Figure 1

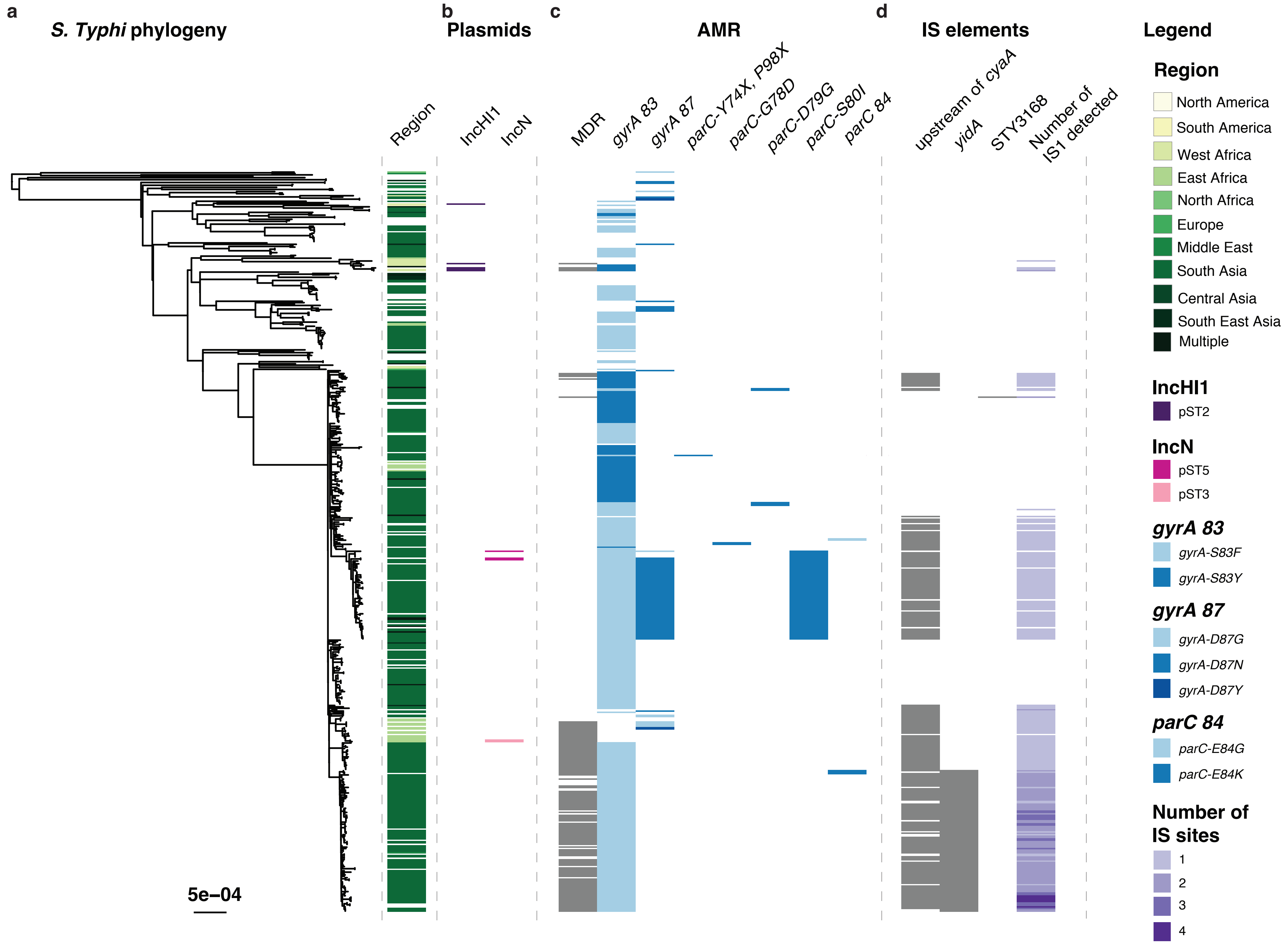
